# Influenza-induced tuft cell expansion alters ILC-mediated inflammation

**DOI:** 10.64898/2026.05.08.723776

**Authors:** Gentile, Michael Maiden, Madeline Singh, Evelyn Martinez, Harshini Kelam, Nicholas Holcomb, Joanna Wong, Meryl Mendoza, Diana Abraham, Alena Klochkova, Sara Kass-Gergi, Andrew Vaughan

**Author notes:** Contributed Equally (Gentile, Michael Maiden).

## Abstract

Tuft cells act as sentinels that amplify type 2 inflammation primarily by activating type 2 innate lymphoid cells (ILC2s). Although normally absent from the distal lung, ectopic tuft cells form after severe lung injury including influenza infection in mice. Here, we investigated the function of these ectopic tuft cells in shaping innate immunity following influenza injury. We observed that IFNγ restrains tuft cell differentiation, whereas ILC2s drive tuft cell expansion, establishing a reciprocal regulatory axis. Tuft cell–deficient mice exhibited reduced eosinophilic inflammation and expansion of ILC1s and ILC3s after influenza injury resolution. Single-cell RNA-seq of influenza infected whole lung revealed transcriptional signatures consistent with type 1 pathway activation, type 2 suppression and oxidative stress. Following influenza injury and subsequent *Alternaria alternata* challenge, tuft cell–deficient mice also showed neutrophilic and ILC3 expansion. Together, these data identify a distal-airway tuft-cell–ILC2 circuit that helps maintain a balanced inflammatory environment in response to viral injury and aeroallergens.

## Introduction

The distal lung bears appreciable capacity for repair: within gas-exchanging alveoli, quiescent alveolar Type 2 can self-renew and differentiate into alveolar Type 1 cells to restore oxygen-exchanging epithelium following mild injury (1, 2). However, following severe injury such as H1N1 influenza A virus infection, intrapulmonary p63 progenitor cells upregulate cytokeratin 5 (Krt5), migrate and proliferate into injured alveoli forming “epithelial scars” that restore barrier function at the likely cost of gas exchange (3–6). These dysplastic regions have been found up to 1-year following injury where mice also showed significantly reduced oxyhemoglobin saturations, demonstrating their long-term impact on lung function (7). Intriguingly, our group and others have identified ectopic tuft cells within these dysplastic regions following IAV infection in mice (8–11).

Historically, tuft cells have held varying names depending on anatomic location including brush cells, microvillus cells and solitary chemosensory cells (12). They are distributed throughout the body including conjunctiva, gingiva, upper airway, thymus, gallbladder, urethra and intestinal tract (13). Tuft cells express components of taste transduction signaling pathways, including bitter taste receptors (T1Rs), sweet and umami taste receptors (T2Rs), Gnat3 (α-gustducin), succinate receptor (SUCNR1) and the transient receptor potential cation channel subfamily M member 5 (TRPM5) to drive chemosensory responses (14–16). Tuft cells also express doublecortin-like kinase 1 (DCLK1) a protein kinase involved in neuronal migration, a marker commonly used for their identification *in situ* (17). The POU homeodomain transcription factor Pou2f3 is necessary for their development (18–20). Regardless of their location, tuft cells function as sentinels that detect pathogens—including viruses, bacteria, aeroallergens and helminths— using taste cell–like chemosensory pathways to orchestrate downstream immune responses (21, 22).

CD4⁺ T lymphocytes, known as helper T cells, are classified into Type 1 helper T cells (T_h_1) and Type 2 helper T cells (T_h_2) based on the cytokines they produce (23). T_h_1-type cytokines are pro-inflammatory and are responsible for intracellular pathogen killing and autoimmune responses (23). T_h_2-type cytokines, including interleukins (IL)-4, IL-5, and IL-13, promote atopy through IgE-mediated eosinophilic responses and anti-inflammatory effects via IL-10 (23, 24). At homeostasis, these two arms of the immune system are balanced (25). Innate lymphoid cells (ILCs) act as counterparts to T_h_1/T_h_2 cells, rapidly shaping the early immune environment (26). ILC2s promote type 2 immune responses via secretion of type 2 cytokines and are counterbalanced by ILC1s which drive type 1 immune responses via the secretion of IFNγ (27, 28). Together, these cells form a reciprocal regulatory axis where the activation of one group of ILCs tilts the overall immune environment toward type 1 or type 2 immunity (29). In parallel, a third immune axis (Type 3) driven by ILC3s promotes steroid-resistant asthma and neutrophilic inflammation through secretion of IL-17 and IL-22 (24).

Tuft cells are well described to form an epithelial-ILC2 positive feedback circuit in the small intestine and upper respiratory tract. In response to helminth infections in the intestine, tuft cells are the sole producers of IL-25, which alongside cysteinyl leukotrienes activates ILC2s to produce IL-13, driving an anti-parasitic immune response and forming a positive feedback loop that drives further tuft cell expansion through an IL-4Rl1l-dependent signaling cascade (30–33). Similarly, in patients with chronic rhinosinusitis with nasal polyps, tuft cells are a major source of IL-25 which drives ILC2 expansion promoting to type 2 inflammation and tuft cell differentiation (34). More recently, rhinovirus infection in immature mice revealed that tuft-cell–derived IL-25 induces ILC2 expansion and promotes type 2 inflammation and tuft cell expansion. Importantly, they also demonstrated tuft cell-dependent airway hyperresponsiveness and mucous metaplasia (35, 36).

Unexpectedly, given these reports, we previously demonstrated that canonical ILC2-derived cytokines IL-4, and IL-13 have no direct role in ectopic tuft cell differentiation or expansion after H1N1 influenza injury (10). Further, we did not observe any obvious effects on epithelial Krt5^+^ dysplasia or goblet cell metaplasia in the absence of tuft cells (10). Here, building on our previous findings, we asked whether ectopic tuft cells contribute to an ILC2 response circuit in the distal airways following influenza infection by systematically disrupting key components of the circuit and assessing changes in tuft cell abundance, immune cell populations and cytokines. We observed that IFNγ restricts tuft cell differentiation, while ILC2s promotes tuft cell expansion. We also found that tuft cell deficient mice (Pou2f3^-/-^) have reduced eosinophils with concomitant expansion of ILC1s and ILC3s, though proportionally fewer ILC2s, following influenza infection. Further, using single-cell RNA-Seq in Pou2f3^-/-^ mice following influenza infection, we found a transcriptomic signature consistent with type 1 pathway activation, type 2 pathway suppression and oxidative stress. Consistent with our transcriptomic data, CXCL5, a neutrophil chemoattractant (37), was increased in bronchoalveolar lavage fluid (BALF) from these mice. Similarly, in Pou2f3^-/-^ mice first infected with influenza, allowed to recover and then challenged with the aeroallergen *Alternaria alternata*, we observed an increase in neutrophils and ILC3s. Taken together, these results support the existence of a distinct tuft cell–ILC2 circuit in the lower airways that serves to balance inflammatory responses to viral infections and aeroallergens.

## Results

### IFNγ limits pulmonary tuft cell numbers and ILC2s following influenza infection

IFNγ represses allergic inflammation and lower levels of IFNγ are associated with stronger type 2 immune responses (38, 39). To determine if tuft cell expansion is influenced by this immune context, we infected IFNγ deficient mice (IFNγ^-/-^) and wildtype mice with influenza (PR8 strain) (Figure 1A). Accordingly, we observed a significant increase in the type 2 cytokine IL-5 in BALF (Figure 1B) and a concomitant increase in lung ILC2 cells at day 9 post PR8-infection in IFNγ^-/-^ versus wildtype mice (Figure 1C).

**Figure 1:**
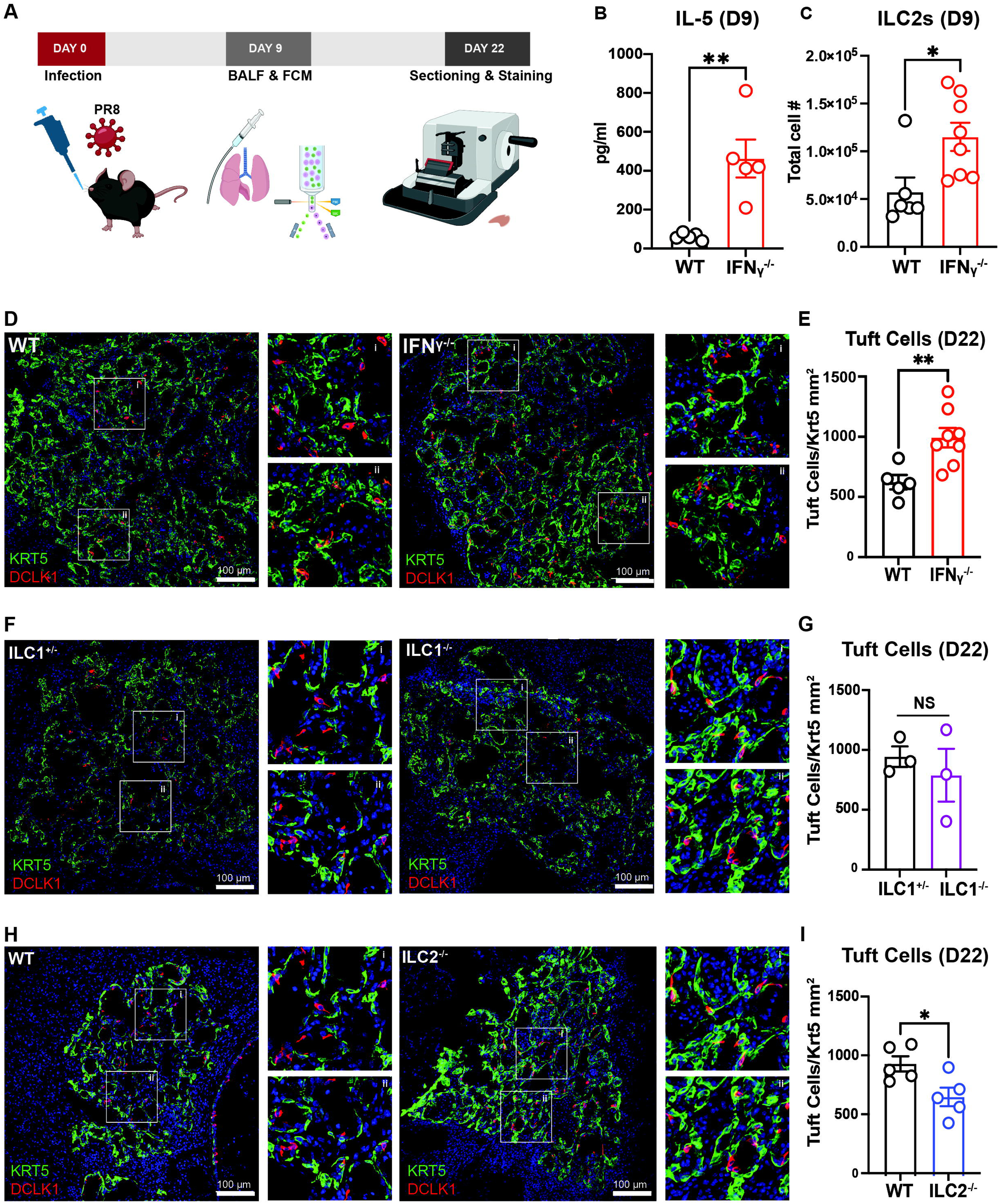
Ectopic pulmonary tuft cells participate in an ILC2-dependent circuit in the lower airways and are restrained by IFNγ after PR8 infection. A) Experimental design outlining intranasal PR8 infection followed by collection of bronchoalveolar lavage fluid (BALF) and lungs for flow cytometry (FCM) at day 9 (D9) post-PR8 infection (p.i.) and immunostaining and tuft cell quantification in the lungs at day 22 (D22) p.i. B) Protein concentration of IL-5 in the BALF of IFNγ ^-/-^mice (n=5) compared to WT (BL6) mice (n=5) at D9 p.i. C) Total cell numbers of type 2 lung innate lymphocytes (ILC2s) (KLRG1^+^NK1.1^-^ of CD127^+^Lineage (Lin)^-^ cells) in IFNγ^-/-^mice (n=8) compared to WT mice (n=6) at D9 p.i. Mice that lost <10% of starting body weight after influenza infection were excluded. D-E) Representative immunostaining and total tuft cell numbers/Krt5+ area in IFNγ ^-/-^ mice (n=8) compared to WT controls (n=5), F-G) in ILC1^-/-^ mice (n=3) compared to ILC1^+/-^ controls (n=3) and in H-I) in ILC2^-/-^ mice (n=5) compared to controls (n=5) at D22 p.i. DAPI is stained in blue, Krt5 in green and Dclk1 in red. Insets show magnified views (i, ii). Scale bars are 100 µm and images are cropped from the 20x z-stack image. H-I) Mice that loss <15% of starting body weight after influenza infection were excluded. B, C, E and I data combined from two-independent experiments. Each circle represents an individual mouse. P values were calculated using unpaired, two-tailed parametric t test (* = p < 0.05, ** = p<0.01, NS = non-significant). Error bars = SEM.

Changes in body weights were calculated as a proxy for inflammation and disease severity finding no significant difference (Supplemental Figure 1A). Pulse oximetry was performed prior to infection and at days-10, 16 and 21 post PR8-infection and there was no difference in oxyhemoglobin saturation between IFNγ^-/-^ and controls (Supplemental Figure 1B). Despite this, we detected a significant increase in total tuft cells present within Krt5^+^ areas in IFNγ^-/-^ lungs compared to controls 22 days following PR8-infection (Figure 1D-E), suggesting that unlike Type I and Type III interferons (10), IFNγ acts to constrain tuft cell expansion. These data suggests that IFNγ acts to limit both tuft cell and ILC2s following influenza infection, similar to its role in regulating the tuft cell–ILC2 circuit in other parts of the body (30–32).

Given that IFNγ secretion by ILC1s can restrain type 2 inflammation which could limit tuft cell expansion (39), we also infected ILC1 deficient mice (*Rroid^-/-^*) with PR8 (40). We found ILC1s were dispensable for tuft cell expansion (Figure 1F-G). Again, we measured changes in body weight and performed pulse oximetry, finding no significant differences between ILC1^-/-^ mice and ILC1^+/-^ controls (Supplemental Figure 1C-D).

### ILC2s contribute to the expansion of pulmonary tuft cells following PR8-infection

We previously demonstrated that unlike the gastrointestinal tract, tuft cell expansion post-influenza is independent of classic ILC2-derived signals like IL-4 and IL-13. To assess whether ILC2s play any remaining role, we infected ILC2 deficient mice (ILC2^-/-^) and wildtype mice with PR8. These transgenic mice lack an enhancer required to drive *Id2* expression (locus control region 1, *LCR1*) in ILC2 precursors, blocking their development (41).Consistent with the literature demonstrating a positive feedback loop between ILC2s and tuft cells (30–32, 34–36), we observed a significant reduction in total tuft cell number per Krt5^+^ area in ILC2^-/-^ lungs compared to controls (Figure 1H-I), but no other obvious changes in overt responses to influenza injury (Supplemental Figure 1E-F). These data suggests that changes in tuft cell number does not impact disease severity or lung damage consistent with our prior findings (10), but that lung ILC2s do provide a signal impacting tuft cell differentiation. Together, these data suggests that ectopic tuft cells form part of a ILC2-dependent circuit in the lower airways.

### Tuft cell deficiency alters ILC and eosinophilic responses to influenza infection

Next, we infected tuft cell deficient mice (Pou2f3^-/-^) and heterozygous controls (Pou2f3^+/-^) with PR8 and used flow cytometry to examine immune response (Figure 2A). We observed no significant differences in body weight or in total lung cell number in Pou2f3^-/-^ mice compared to controls at D22 post PR8-infection (Supplemental Figure 2A-B).

**Figure 2:**
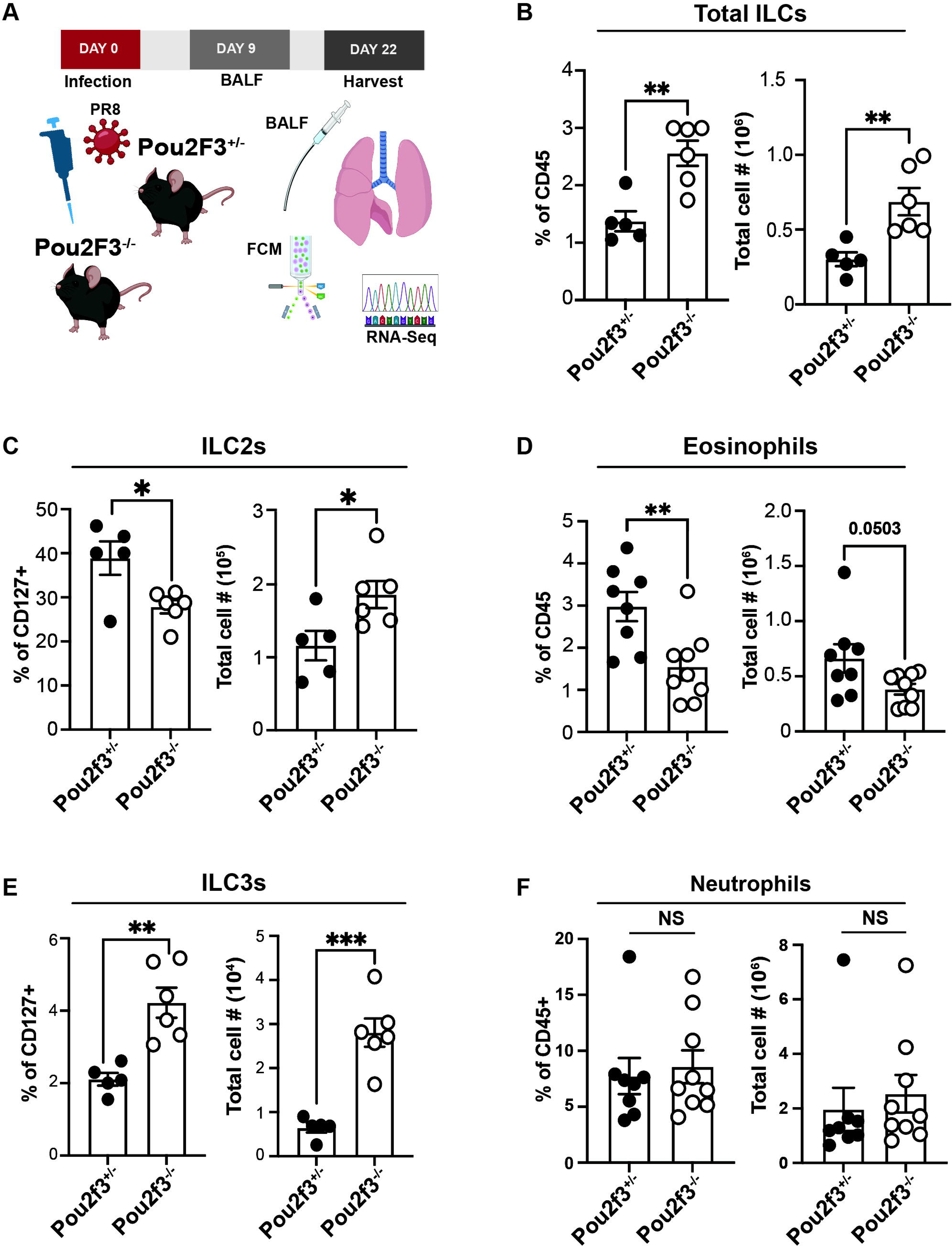
Tuft cell–deficient mice have reduced lung eosinophils and increased ILCs following influenza infection. **A**) Experimental design outlining intranasal PR8 infection in Pou2f3^+/-^ and Pou2f3^-/-^ mice followed by collection of bronchoalveolar lavage fluid (BALF) at D9 p.i. and lungs for flow cytometry analysis as well as single-cell RNA sequencing (RNA-Seq) at D22 p.i. **B**) Frequency of CD45^+^ and total cell numbers of innate lymphocytes (ILCs) (CD127^+^ of Lin^-^CD11b^low/int^ cells) in the lungs of Pou2f3^+/-^ and Pou2f3^-/-^ mice at D22 p.i. **C**) Frequency of CD127^+^ and total cell numbers of ILC2s (KLRG1^+^NK1.1^-^ of CD127^+^ Lin^-^cells) and **D**) frequency of CD45^+^ and total cell numbers of eosinophils (CD11c^low/int^ MHCII^-^ of SiglecF^+^F4/80^+^ cells) in the lungs of Pou2f3^+/-^ and Pou2f3^-/-^ mice at D22 p.i. **E)** Frequency of CD127^+^ and total cell numbers of ILC3s (Rorgt^+^Tbet^-^ of KLRG1^-^CD127^+^cells) and **F**) frequency of CD45^+^ and total cell numbers of neutrophils (Ly6g^+^CD11b^+^ of live CD45^+^ cells) in the lungs of Pou2f3^+/-^ and Pou2f3^-/-^ mice at D22 p.i. **D** and **F** data combined from two-independent experiments. Each circle represents an individual mouse. P values were calculated using unpaired, two-tailed parametric t test (* = p < 0.05, ** = p<0.01, *** p < 0.001, NS = non-significant). Error bars = SEM. Full gating strategy for the different immune cell populations found in material and methods.

When investigating the innate immune cell compartment, we observed an overall increase in ILCs as a percentage of immune cells (CD45^+^) in tuft cell deficient mice (Figure 2B). There was also an increase in the absolute number of ILC1 cells compared to controls (Supplemental Figure 2C). Consistent with the hypothesis that tuft cells drive an ILC2 circuit in the distal airways, there was a significant decrease in ILC2 cells as a percentage of CD127^+^ cells in Pou2f3^-/-^ mice compared to controls (Figure 2C and Supplemental Figure 2D). However, unexpectedly, the absolute number of ILC2 cells in Pou2f3^-/-^ mice were greater than controls (Figure 2C), seemingly due to expansion of the overall ILC pool (Figure 2B). Further we observed a significant decrease in eosinophils as a percentage of CD45^+^ cells and a strong decreasing trend in absolute number in Pou2f3^-/-^ mice compared to controls (Figure 2D and Supplemental Figure 2E).

Finally, we saw an increase in ILC3s (Figure 2E), which are known to drive neutrophil production, recruitment and activation (42, 43), though neutrophil numbers themselves did not differ at this time point (Figure 2F). We also observed no differences in alveolar macrophages, inflammatory monocytes or natural killer (NK) cells (Supplemental Figure 2F-H). Together, these data suggest that the presence of tuft cells during viral injury suppresses some components of type 1 and 3 responses, especially as manifested by ILC expansion.

### Transcriptional profiling of immune and non-immune cell types from tuft cell deficient mice

Given that we observed limited changes in inflammation and no change in disease severity as measured by weight loss (Supplemental Figure 2A) after PR8-infection in Pou2f3^-/-^ mice, we pursued an unbiased approach using single cell RNA sequencing to determine what downstream pathways may be regulated by tuft cells. We profiled both uninfected and post-influenza mice on D22 following infection, and separated lung cells from these mice into immune (CD45⁺) and non-immune (CD45⁻) fractions to better resolve the contributions of different cellular populations (Figure 2A).

In non-immune cell types, UMAP visualization revealed global differences in gene expression between Pou2f3^-/-^ mice and controls with transcriptional differences most apparent in general capillaries (gCaps) and alveolar fibroblast (Alv_Fib) (Figure 3A). At day 22 following PR8 influenza infection, 848 genes were differentially expressed in Pou2f3^⁻/⁻^ mice compared with control animals. (adjusted p-value <0.05) (Figure 3B-C). There were a total of 13 genes upregulated, and 14 genes downregulated in Pou2f3^-/-^ mice regardless of infection status. The top 20 differential expressed genes (DEG) for epithelial, endothelial and mesenchymal along with marker genes cells are shown in Figure 3D-F. The transcriptomic profile was consistent with type 1 immune pathway activation, including differential expression of *Retnla*, which suppresses type 2 immunity in epithelial cells (44); *Cxcl15*, a neutrophil-recruiting chemokine expressed in mesenchymal cells; and *Cyp1a1*, which mitigates oxidative stress (45). *Atox1* was also differentially expressed across all cell types and is known to prevent oxidative damage (46). We also saw differential expression of *Socs3* which shifts macrophages toward pro-inflammatory phenotypes and limits IL-10 anti-inflammatory signaling (47).

**Figure 3:**
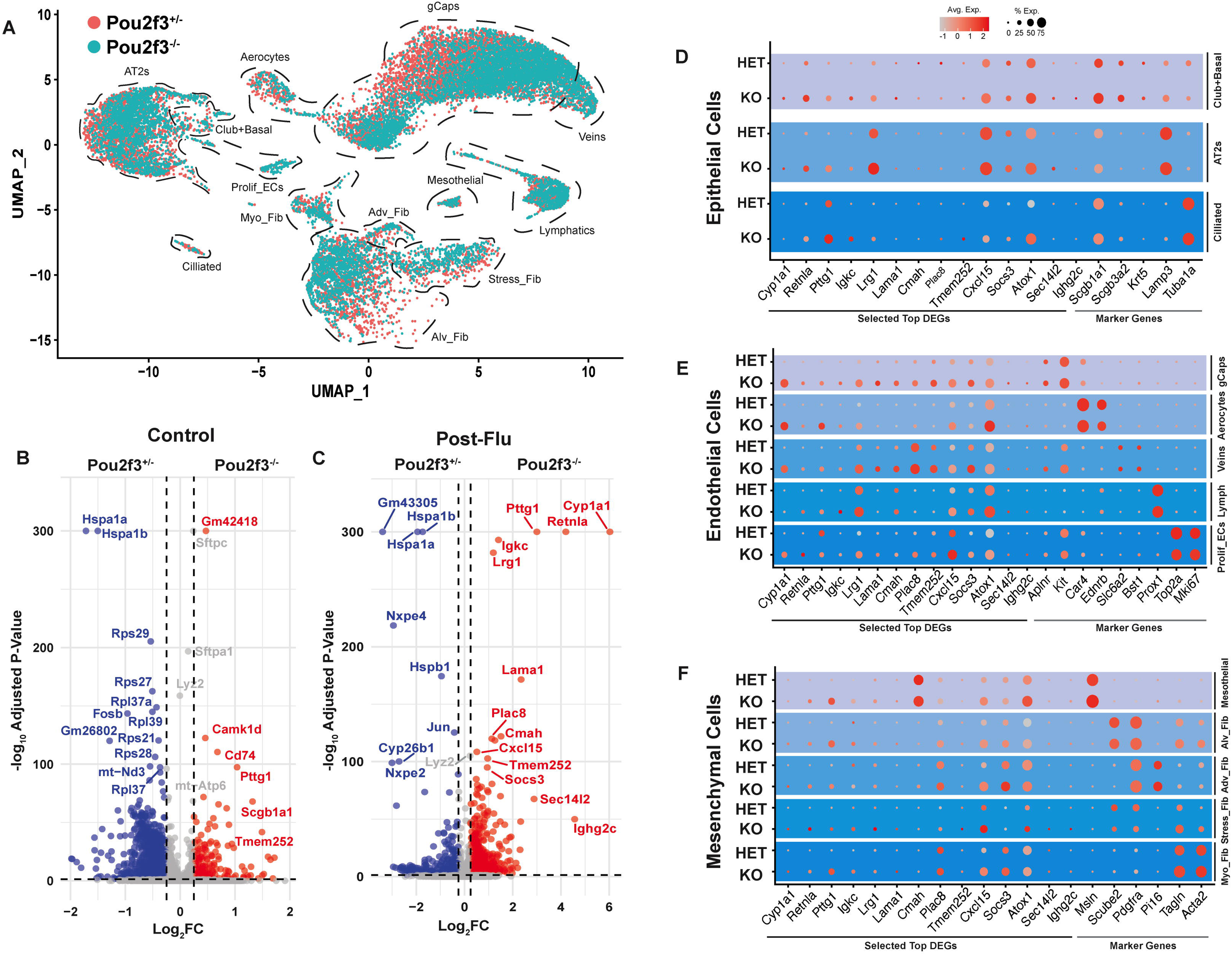
Transcriptional profiling of non-immune cell populations in tuft cell-deficient mice post-influenza. **A**) Single-cell RNA-seq UMAP clustering of sorted non-immune (CD45^-^) cell types from mice at D22 p.i. **B**) Volcano plot comparing gene expression in uninfected Pou2f3^+/-^ and Pou2f3^-/-^ mice. **C**) Volcano plot comparing gene expression in infected Pou2f3^+/-^ and Pou2f3^-/-^ mice. Top 14 differentially expressed genes (DEGs) and selected marker genes in **D**) epithelial, **E)** endothelial, and **F)** mesenchymal cells from Pou2f3^+/−^ and Pou2f3^−/−^ mice at D22 p.i.

Overall, immune cell types exhibited fewer transcriptomic differences compared to non-immune cells (Supplemental Figure 3). We observed differential expression of 556 genes between Pou2f3^-/-^ and controls at D22 post PR8-infection (adjusted p-value <0.05) (Supplemental Figure 3B-C). In immune cell types, there were 9 genes upregulated and 15 genes downregulated in Pou2f3^-/-^ mice, independent of infection status. The observed differential expression was primarily restricted to mitochondrial and housekeeping genes which may indicate apoptosis or changes in metabolic states. (e.g., *mt-Co3, mt-Atp6, mt-Co2, mt-Cytb, mt-Nd* genes, *Ubc, Rps29*) (Supplemental Figure 3D-E). Notably, Pou2f3^-/-^ cells showed consistently higher mitochondrial gene expression, particularly in T and NK cells, suggesting that the Pou2f3 genotype may alter cellular metabolic or activation states without affecting lymphocyte identity.

To further explore differences in inflammatory response as a function of tuft cells, we also performed a Luminex® cytokine array on BALF collected from Pou2f3^-/-^ mice and controls at D9 and D22 post PR8-infection (Figure 2A and Supplemental Figure 4).Supporting our transcriptomic data, we observed an increase in the neutrophil chemoattractant CXCL5/LIX (Supplemental Figure 4K) in Pou2f3^-/-^ mice only, recapitulating recent demonstrations that that tuft cell deficient animals have increased CXCL5/LIX expression and neutrophilia in the gall bladder (48). Otherwise, we found few differences in cytokine profiling in Pou2f3^-/-^ mice compared to controls (Supplemental Figure 4A-M). Taken together, these findings suggest an unanticipated role for tuft cells in influencing a type 1-and 3 associated responses, including the potential to recruit neutrophil and modulating ILC expansion during recovery from influenza injury.

### Tuft cell deficient mice have an increased pulmonary neutrophilic response following *Alternaria* challenge

As previously noted, Pou2f3^-/-^ mice infected with PR8 had a decreased frequency of ILC2s and eosinophils (Figure 2C and D). Given their established role in type 2 inflammation in the upper airways (13, 34–36), we hypothesized that tuft cells may promote a pro-asthmatic immune milieu in the distal airways following influenza infection. To test this, we used a two-hit model in which mice were first infected with PR8, allowed to recover for 3-weeks, and then challenged with the aeroallergen *Alternaria alternata* followed by flow cytometry to assess the inflammatory contribution of tuft cells (49) (Figure 4A).

**Figure 4:**
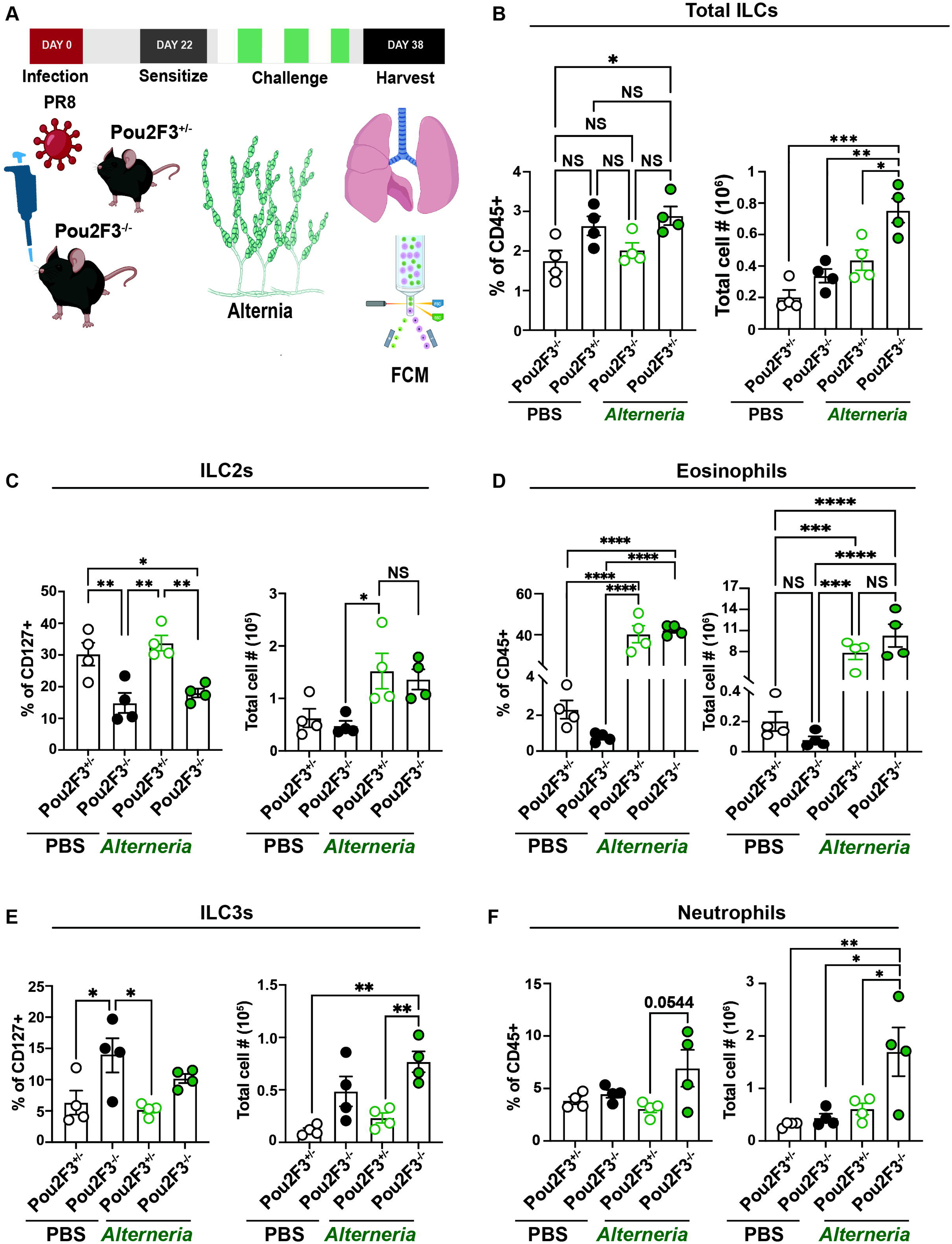
Pulmonary neutrophil and ILC responses are elevated in tuft cell–deficient mice after *Alternaria alternata* secondary challenge. **A)** Experimental design outlining intranasal PR8 infection followed by allergen sensitization (40 µg) and challenge (20 µg) as well as collection of lungs for flow cytometry analysis in Pou2f3^+/-^ and Pou2f3^-/-^ mice at D38 p.i. **B)** Frequency of CD45^+^ and total cell numbers of ILCs (CD127^+^ of Lin^-^CD11b^low/int^ cells) in *Alternaria alternata* or PBS challenged Pou2f3^+/−^ and Pou2f3^-/−^ mice at D38 p.i. **C**) Frequency of CD127^+^ cells and total cell numbers of ILC2s (KLRG1^+^NK1.1^-^ of CD127^+^ Lin^-^ cells) and **D**) frequency of CD45^+^ cells and total cell numbers of eosinophils (CD11c^low/int^ MHCII^-^ of SiglecF^+^CD11B^int/high^ cells) in the lungs of *A. alternata* or PBS challenged Pou2f3^+/-^ and Pou2f3^-/−^ mice at D38 p.i. **E**) Frequency of CD127^+^ cells and total cell numbers of ILC3s (Rorgt^+^ of KLRG1^-^NK1.1^-^CD127^+^ cells) and **F**) frequency of CD45^+^ cells and total cell numbers of neutrophils (Ly6g^+^CD11b^+^ of live CD45^+^ cells) in the lungs of *A. alternata* or PBS challenged Pou2f3^+/-^ and Pou2f3^-/−^ mice at D38 p.i. Each circle represents an individual mouse. P values were calculated using one-way ANOVA with Tukey’s post-test for multiple comparisons (* p < 0.05, ** p < 0.01, *** p < 0.001, **** p< 0.0001. NS = non-significant). Error bars = SEM. Complete gating strategy for the different immune cell populations found in material and methods.

We observed no change in body weight throughout the course of the experiment (Supplemental Figure 5A). However, we did observe an increase in total cell numbers in the lung upon *A. alternata* challenge regardless of genotype (Supplemental Figure 5B). At day 37 following PR8 infection, pulse oximetry measurements revealed a significant decrease in oxyhemoglobin concentrations in Pou2f3^+/-^ control mice exposed to *A. alternata*, consistent with impaired gas exchange (Supplemental Figure 5C). In contrast, Pou2f3^-/-^mice showed no significant change in oxyhemoglobin saturations. These data suggest that tuft cells may play a role in suppressing airway inflammation and dysfunction following aeroallergen challenge.

Next, we assessed the impact of Pou2f3 deletion on innate immune cell populations. We observed a significant increase in total ILCs in Pou2f3^-/-^mice challenged with *A. alternata* compared to controls (Figure 4B). There was also an increase in the absolute number of ILC1s in mice challenged with *A. alternata* relative to PBS challenge, regardless of genotype (Supplemental Figure 6A). In keeping with a role for tuft cells in an epithelial-ILC2 circuit, Pou2f3^-/-^ mice had decreased frequencies of pulmonary ILC2s relative to Pou2f3^+/-^ controls (Figure 4C). There was no significant change in total cell numbers (Figure 4C), likely due to expansion of the overall ILC pool (Figure 4B). Surprisingly, there was no difference in pulmonary eosinophils between Pou2f3^-/-^ and Pou2f3^+/-^ mice challenged with *A. alternata* at D38 following PR8 infection (Figure 4D).

Analysis of ILC3 populations, which drive neutrophilic inflammation (42, 43), following *A. alternata* challenge revealed significantly increased total numbers in Pou2f3^-/-^ mice compared to Pou2f3^+/-^ controls (Figure 4E). Intriguingly, there was a significant increase in lung neutrophils in Pou2f3^-/-^ mice challenged with *A. alternata* compared to Pou2f3^+/-^ mice (Figure 4F). This is consistent with our prior observation that there was an increase in the neutrophil chemoattractant CXCL5/LIX in the BALF of Pou2f3^-/-^ mice at D22 post PR8-infection (Supplemental Figure 4K). We also found no differences in additional immune cell types including alveolar macrophages, inflammatory monocytes, or NK cells between Pou2f3^-/-^ and Pou2f3^+/-^ mice challenged with *A. alternata* (Supplemental Figure 6B-D). Taken together, these data suggest that tuft cells partially constrain neutrophilic inflammation following aeroallergen challenge.

## Discussion

Although tuft cells are critical chemosensory regulators of immunity to helminths and bacteria, their responses to viral infections and aeroallergens remain poorly understood, especially in the context of the distal airways (13). In the upper respiratory tract and intestine, tuft cells regulate an epithelial-ILC2 circuit that drives type 2 immunity that is dependent on both IL-25 and Il-4rl1l signaling (30–32). Conversely, our previous work demonstrated that post-IAV ectopic tuft cells develop independently of IL-25 and IL-4Rl1l signaling (10) but still rely upon the master transcription factor POU2F3 for their development (10) and share transcriptomic features with both intestinal and tracheal tuft cells, suggesting analogous functional roles (21). To explore these possible functions as well as what upstream signals are required for tuft cell development, we genetically disrupted key components of the ILC2-circuit and assessed their relative contributions to circuit function.

Our findings generally reinforce a role for tuft cells within an epithelial–ILC2 regulatory circuit in the lower airways, albeit with some interesting differences compared to existing models. IFNγ can suppresses ILC2 cytokine production (29, 50), and accordingly, we found that IFNγ restricts both ILC2 and tuft cell expansion (Figure 1C, D and E). In contrast, ILC2s promoted post-influenza tuft cell expansion (Figure 1H and I), suggesting a reciprocal regulatory axis between tuft cells and ILC2s. While we presume the effects of IFNγ are acting largely through inhibition of ILC2 expansion (39), further work is required to demonstrate this definitively. Further, the identity of the ILC2-derived signal promoting tuft cell differentiation remains unknown, an especially intriguing open question given previous data suggesting neither IL-4 or IL-13 are critical for their development post-influenza.

We did not observe an obviously attenuated injury or inflammatory response in tuft cell deficient mice following influenza infection, likely due to the emergence of ectopic tuft cells in the distal airways during the recovery phase. Upon recovery from influenza infection, tuft cell–deficient mice exhibited a reduction in frequency of lung eosinophils and ILC2s (Figure 2C-D). We also observed an expansion of ILC1s and ILC3s (Supplemental Figure 2C and Figure 2E), as well as increased CXCL5/LIX in BALF (Supplemental Figure 4K) at D22 post PR8-infection. In keeping with these observations, a variety of gene expression changes observed in single-cell RNA-seq further suggested activation, or at least poised activation, of type 1 inflammation-associated pathways in tuft cell–deficient mice following influenza infection. Therefore, we challenged Pou2f3^-/-^ mice with the fungi *A. alternata* to see if they would have a decreased asthmatic phenotype. We did not observe a significant decrease in eosinophil or ILC2 numbers in tuft cell deficient mice challenged with *A. alternata* compared to controls (Figure 4C-D). Instead, we observed an increase in neutrophils and ILC3 numbers (Figure 4E-F).

Though the lack of tuft cell-dependent changes in eosinophilia was somewhat surprising, we predict that the prolonged sensitization to *A. alternata* extract over several weeks may result in antigen-specific T_h_2 T cell responses overriding the influences of more acute innate Type 2 responses promoted by tuft cells and ILC2s, a possibility suggesting future study. Still, overall these results support the existence of a tuft cell–ILC2 regulatory loop in the distal lung that may help to maintain immunologic equipoise between type 1 and 2 inflammatory responses.

Previously, we found that tuft cells are not required for the formation of Krt5^+^ cells, goblet cells, or conversion of Krt5^+^ cells to AT2s after severe influenza injury (10), though this does not preclude the possibility of tuft cells contributing to differential inflammatory responses or other less obvious effects. Here, we found that ectopic tuft cells form part of an epithelial-ILC circuit in the lower airways and limit neutrophilic inflammation following viral infection and aeroallergen challenge. These findings highlight unexpected differences and complexities in tuft cell signaling in the lung compared to more well studied tissues like the intestines. Future characterization of these signals may offer additional insights into the enigmatic functions of these intriguing cells.

## Methods

### Animals

C57BL6/J, *IFN*γ^-/-^, ILC1^+/-^, ILC1^-/-^, ILC2^-/-^ (41), *POU2F3*^+/-^, and *POU2F3*^-/-^ (19) mice were used. Adult mice of both sexes were used and all mice are on a C57BL6/J background unless otherwise noted. The protocol number associated with the ethical approval of this work is 806262 (University of Pennsylvania). For all animal studies, no statistical method was used to predetermine sample size. The experiments were not randomized, and the investigators were not blinded to allocation during experiments and outcome assessment.

### IAV infection and Aeroallergen Challenge

Infections were performed as previously described (10). All viral infections utilized influenza strain A/H1N1/PR/8 obtained from Dr. Carolina Lopez (51). For influenza infection at the University of Pennsylvania, virus was administered intranasally. Mice ranging between 15 and 20 g in weight were infected with 30 tissue culture infectious dose (TCID) 50 units of PR8, mice weighing between 20 and 25 g were given 40 TCID50 units, and mice ranging between 25 and 30 g in weight were given 50 TCID50 units. For aeroallergen experiments, mice were first sensitized with 40 µg of *Alternaria alternata* extract intranasally. They were then challenged with 6 doses of *A. alternata* intranasally (20 µg) to induce allergen sensitivity as previously described (41, 49). Control mice received Phosphate-Buffered Saline (PBS). PR8-infected mice that did not survive to the end point of the experiment or did not lose weight were excluded.

### Whole-lung single-cell suspension preparations

Lungs were harvested from mice and single-cell suspensions were prepared. Briefly, lungs were perfused with ice cold PBS via the left atrium, lobes were separated and collected in ice cold Dulbecco’s Modified Eagle Medium (DMEM; Thermo Fisher Scientific) +10% fetal bovine serum (FBS; Thermo Fisher Scientific). Once all mice were harvested, the lobes were placed in 1.5 mL of digestive media (DMEM with 0.15 mg/ml of Liberase TM (Roche # 5401119001), 0.014 mg/mL of DNase I (Sigma Aldrich # D4527-20KU) and 0.20 mg/mL of Dispase II (Sigma Aldrich D4693-1G), cut into small pieces and placed at 37°C to shake for 40-mins. The samples where then passed through a 16-G needle three times and placed back shaking at 37°C for 15-mins. The samples were then passed through a 18-G needle three times and filtered through a 70 µm strainer. The filters were washed with DMEM+10% FBS. Red blood cells were then lysed for 3-mins (Thermo Fisher Scientific #A1049201) and cells were manually counted using a hemacytometer. Whole-lung single-cell suspensions were then used for subsequent experiments.

### Flow cytometry for immunophenotyping

Cell suspensions were washed twice in PBS and then incubated with a fixable viability dye (eFluor 780 eBioscience # 65-0865-18) for 20-mins at 4°C. Afterwards, the cells were incubated with TruStain FcX (anti-mouse CD16/32) Antibody (BioLegend, #101319) for 10-mins at 4°C. The cells were then stained with the following antibodies in appropriate combinations of fluorophores. B220 (clone: RA3-6B2), CD3 (clone: 145-2C11), CD11b (clone: M1/70), CD11c (clone: N418), CD45 (clone: 30-F11),), CD127 (clone: A7R34), F4/80 (clone: BM8), KLRG1 (clone: 2F1/KLRG1), Ly6c (clone: HK1.4), Ly6g (clone: 1A8), MHCII (clone: M5/114.15.2), NK1.1 (clone: PK136), Rorgt (clone: Q31-378), Siglec-F (clone: E50-2440), and Tbet (clone: 4B10). The samples were then fixed with 3.2% paraformaldehyde for 12-mins at 4°C. For intracellular staining of transcription factors, cells were fixed and permeabilized with the FoxP3 Fix/Perm kit (eBiosciences) according to the manufacturer’s instructions. Data were acquired with a FACSymphony A3 (BD Biosciences) and analyzed using FlowJo software.

Gating strategy for Figure 1C, Figure 2 and Supplemental Figure 2 defining: Alveolar Macrophages (singlet Live CD45^+^Ly6g^-^ Ly6c^low/int^ F4/80^+^SiglecF^+^MHCII^+^CD11c^+^), Eosinophils (singlet Live CD45^+^Ly6g^-^ Ly6c^low/int^ F4/80^+^SiglecF^+^MHCII^-^CD11c^low/int^), ILCs (singlet Live CD45^+^Lin^-^ (CD3^-^B220^-^) CD11b^int/low^CD127^+^), ILC1s (singlet Live CD45^+^Lin^-^CD11b^int/low^CD127^+^KLRG1^-^Rorgt^-^Tbet^+^), ILC2s (singlet Live CD45^+^Lin^-^CD11b^int/low^CD127^+^NK1.1^-^KLRG1^+^), ILC3s (singlet Live CD45^+^Lin^-^CD11b^int/low^CD127^+^KLRG1^-^ Tbet^-^ Rorgt^+^), Inflammatory Monocytes (singlet Live CD45^+^Ly6g^-^CD11b^+^Ly6c^high^), Natural Killer cells (singlet Live CD45^+^Lin^-^CD11b^int/low^CD127^-^NK1.1^+^) and Neutrophils (singlet Live CD45^+^CD11b^+^Ly6g^+^).

Gating strategy for Figure 4 and Supplemental Figure 6 defining Alveolar Macrophages (singlet Live CD45^+^Ly6g^-^SiglecF^+^CD11b^int/high^MHCII^+^CD11c^+^), Eosinophils (singlet Live CD45^+^Ly6g^-^SiglecF^+^CD11b^int/high^ MHCII^-^CD11c^low/int^), ILCs (singlet Live CD45^+^Lin^-^ (CD3^-^B220^-^) CD11b^int/low^CD127^+^), ILC1s (singlet Live CD45^+^Lin^-^CD11b^int/low^CD127^+^KLRG1^-^NK1.1^+^), ILC2s (singlet Live CD45^+^Lin^-^CD11b^int/low^CD127^+^NK1.1^-^KLRG1^+^), ILC3s (singlet Live CD45^+^Lin^-^CD11b^int/low^CD127^+^KLRG1^-^ NK1.1^-^ Rorgt^+^), Inflammatory Monocytes (singlet Live CD45^+^Ly6g^-^SiglecF^-^ CD11b^+^Ly6c^high^), Natural Killer cells (singlet Live CD45^+^Lin^-^ CD11b^int/low^CD127^-^NK1.1^+^) and Neutrophils (singlet Live CD45^+^CD11b^+^Ly6g^+^).

### Fluorescence-activated cell sorting for single cell RNA-Seq

Whole-lung single-cell suspensions were prepared as above and as previously described (10). First, Whole-lung single-cell suspensions were pelleted for 5-mins at 550xg at 4°C. Pellets were re-suspended in in sort buffer (SB; DMEM+2% Cosmic Calf serum (CC; Thermo Fisher Scientific) + 1% penicillin and streptomycin). Cells were blocked with 1:50 TruStain FcX (anti-mouse CD16/32 antibody; BioLegend, #101319) for 10-mins at 37°C. For immune cells we performed EasyStep^TM^ Mouse CD45 Positive Selection Kit (StemCell Technologies) per manufacture’s protocol. For nonimmune cells, the cell suspension was stained using allophycocyanin/Cy7--conjugated rat anti-mouse CD45 antibody (1:200, BioLegend, #101319), PE-conjugated rat anti-mouse EpCam antibody (1:500, BioLegend, G8.8, #118206). Stained cells and ‘fluorescence minus one’ controls were then resuspended in SB + 1:1000 Dnase + 1:1000 Draq7 (BioLegend, #424001) as a live/dead stain. Nonimmune cells were isolated as all live CD45^neg^ cells. All FACS sorting was done on a BD FACS AriaFUSION Sorter (BD Biosciences).

### Single-cell RNA-seq

Whole-lung single-cell suspensions were prepared as above and as previously described (10). Single-cell RNA-Seq was performed using the Chromium System (10x Genomics) and the Chromium Single Cell 3’ Reagent Kits v2 (10x Genomics) at the Children’s Hospital of Philadelphia Center for Applied Genomics. After sequencing, initial data processing was performed using Cellranger (v.3.1.0). Cellranger mkfastq was used to generate demultiplexed FASTQ files from the raw sequencing data. Next, Cellranger count was used to align sequencing reads to the mouse reference genome (GRCm38) and generate single-cell gene barcode matrices. Post-processing and secondary analysis were performed using the Seurat package (v.4.4.1). First, variable features across single cells in the dataset will be identified by mean expression and dispersion. Identified variable features was then be used to perform a principal component analysis (PCA). The dimensionally reduced data was used to cluster cells and visualize using a Uniform Manifold Approximation and Projection (UMAP) plot.

### Tissue preparation for immunofluorescence

Each lung was prepared as previously described (10). Briefly, lungs were perfused with PBS via the left atrium and then perfused with ice-cold 3.2% paraformaldehyde (PFA; Thermo Fisher Scientific) and placed in a 50 mL tube with 25 mL of PFA to shake at room temperature (RT) for 1-hr. Following incubation, the lungs were washed twice in PBS for 1-hr at RT. Lungs were then placed in 30% sucrose (Sigma-Aldrich) overnight shaking at 4°C. The following day, the tissues were placed in 15% sucrose-50% optimal cutting temperature compound (OCT; Fisher Healthcare) shaking for 2-hr at RT. The fixed lungs were then embedded in OCT and flash frozen. Using a cryostat, the lungs were sectioned (6 µm) and stored at −20°C. Lung sections were fixed in 4% PFA for 5-mins at RT and then washed three times with PBS for 5-mins at RT. Slides were blocked for 1-hr at RT in a humid chamber with blocking buffer: 1% BSA, Gold Bio, 5% donkey serum (Sigma), 0.1% Triton X-100 (Fisher BioRe-agents) and 0.02% sodium azide (Sigma-Aldrich) in PBS. The slides were then stained in blocking buffer overnight at 4°C with a combination of primary antibodies. The following day, the slides were rinsed 3 times for 2-mins with PBS +0.1% Tween (Sigma-Aldrich) and then stained with secondary antibodies in blocking buffer for 90-mins at RT in the humidity chamber. The slides were rinsed 3 times for 2-mins with PBS+0.1% Tween and stained with DAPI (1:10,000 dilution; catalog no. D21490, Thermo Fisher Scientific). Slides were then mounted with Fluoroshield (Sigma) and imaged using a Leica inverted fluorescent microscope Dmi8 and analyzed using Las X software. Primary antibodies used: rabbit anti-Dclk1 (1:500 dilution; catalog no. ab37994 or ab31704, Abcam), chicken anti-Krt5 (1:1000 dilution; catalog no. 905901, BioLegend). Secondary antibodies used: donkey anti-rabbit AF568 (1:1000 dilution; catalog no. A10042, Thermo Fisher Scientific, donkey anti-chicken AF488 (1:1000 dilution; catalog no. 703-545-155, Jackson ImmunoResearch)

### Pulse oximetry

Peripheral oxygen saturation (SpO2) was determine as previously described (7). Briefly, using a MouseOx Plus Rat & Mouse Pulse Oximeter and a MouseOx small collar sensor (Starr Life Sciences Corp.) SpO2 what taken at the indicated time points for each experiment. At least one day prior to the initial reading, the mouse’s neck and shoulder area was shaved with a razor at the site where the collar sensor was placed. Recordings were taken using MouseOx Premium Software (Starr Life Sciences Corp., Oakmont, PA, USA). Only measurements with zero error codes were used in our analysis. The overall average of the readings were used to determine the SpO2 for each mouse at the indicated time points.

### Image quantification

Lung sections were imaged on a Leica Dmi8 microscope using a 20×0.75 NA objective. Image quantification was performed by manual area measurements utilizing ImageJ/FIJI for Krt5 area and manual quantification of Dclk1+ cells. We counted total tuft cell number in >/= 2 discrete Krt5^+^ cell area in total lung area by scanning whole lobe sections to ensure adequate representation.

### Chemokine/Cytokine Screen

Cytokines and chemokines were measured using a multiplex bead-based immunoassay (Luminex® xMAP technology) per manufactures instructions (52). Briefly, bronchoalveolar lavage fluid (BALF) was collected from mice using ice-cold PBS (1 mL) as previously described (5). Samples were incubated with antibody-conjugated fluorescent microspheres, followed by biotinylated detection antibodies and streptavidin-phycoerythrin. Beads were analyzed on a Luminex® platform, and analyte concentrations were calculated from standard curves using five-parameter logistic regression performed by the Human Immunology Core at the University of Pennsylvania.

## Statistics

All statistical calculations were performed using GraphPad Prism. Relevant statistical tests utilized are stated in the figure legends.

## Acknowledgements

We thank all Vaughan Lab members for helpful discussions and suggestions. We thank the CHOP Flow Cytometry Core and Center for Human Genomics. We would like to thank Dr. Jorge Henao-Mejia for providing us with the ILC1^-/-^ and ILC2^-/-^ mice. We would like to thank Dr. Boris Striepen for providing the IFNγ^-/-^ mice. We would like to thank Honghong Sun of the Human Immunology Core for her expert technical assistance. Funding: This work was supported by NIH grants RO1HL131817, RO1HL153539 Grant to AEV, T32HL160493 to MMM, and Fonds de recherche du Québec Santé Postdoctoral Fellowship to MEG. Competing interests: The authors declare that they have no competing interests. Data and materials availability: All data needed to the conclusions in the paper are present in the paper and/or the Supplementary Materials. Additional data related to this paper may be requested from the authors.

## Funding

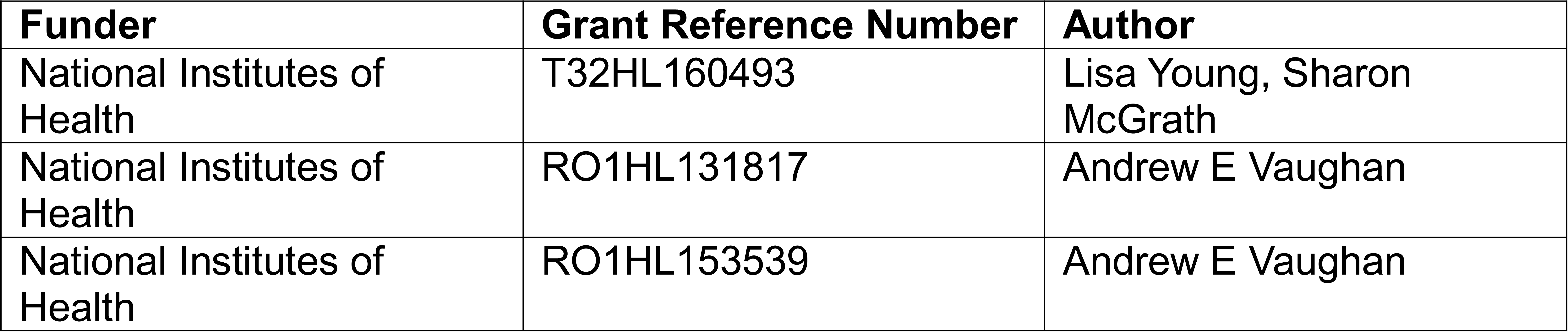

## Author contributions

Michael Maiden, Maria Elena Gentile, Conceptualization, Data curation, Formal analysis, Investigation, Methodology, Writing - original draft, Writing - review and editing; Evelyn Martinez, Data curation; Madeline Singh, Data curation; Harshini Kalem, Data curation; Alena Klochkova, Sara Kass-Gergi, Joana Wong, Nicolas P Holocomb and Diana Abraham, Writing - review and editing; Andrew E Vaughan, Conceptualization, Resources, Data curation, Formal analysis, Supervision, Funding acquisition, Investigation, Methodology, Writing - original draft, Project administration, Writing - review and editing

## Ethics

All animal procedures were approved by the Institutional Animal Care and Use Committee (IACUC) of the University of Pennsylvania. All experiments were performed with every effort to minimize suffering. The protocol number associated with the ethical approval of this work is 806262 (University of Pennsylvania).

