## Supplementary figures and images for "Influenza-induced tuft cell expansion alters ILC-mediated inflammation"

### Supplemental Figure 1

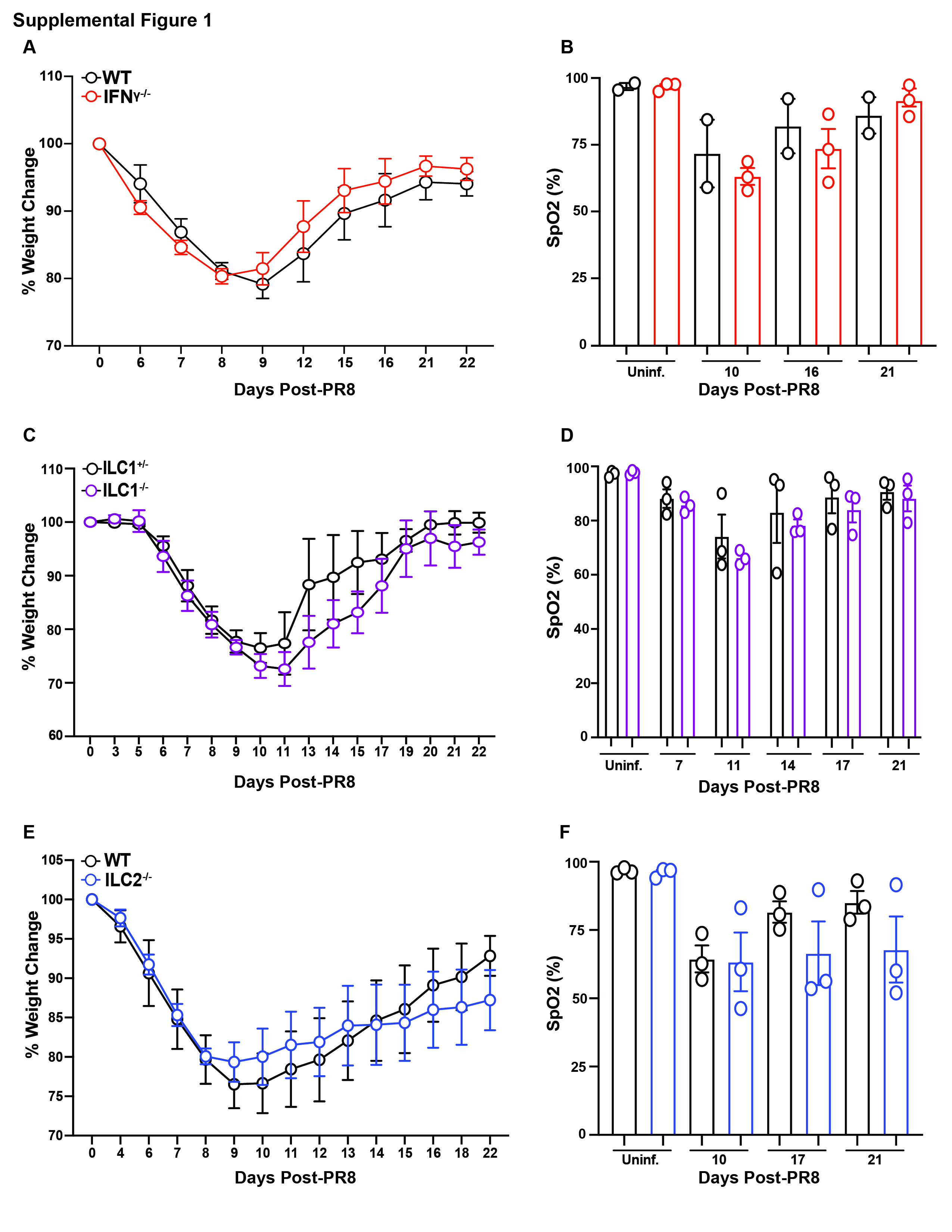

### Supplemental Figure 2

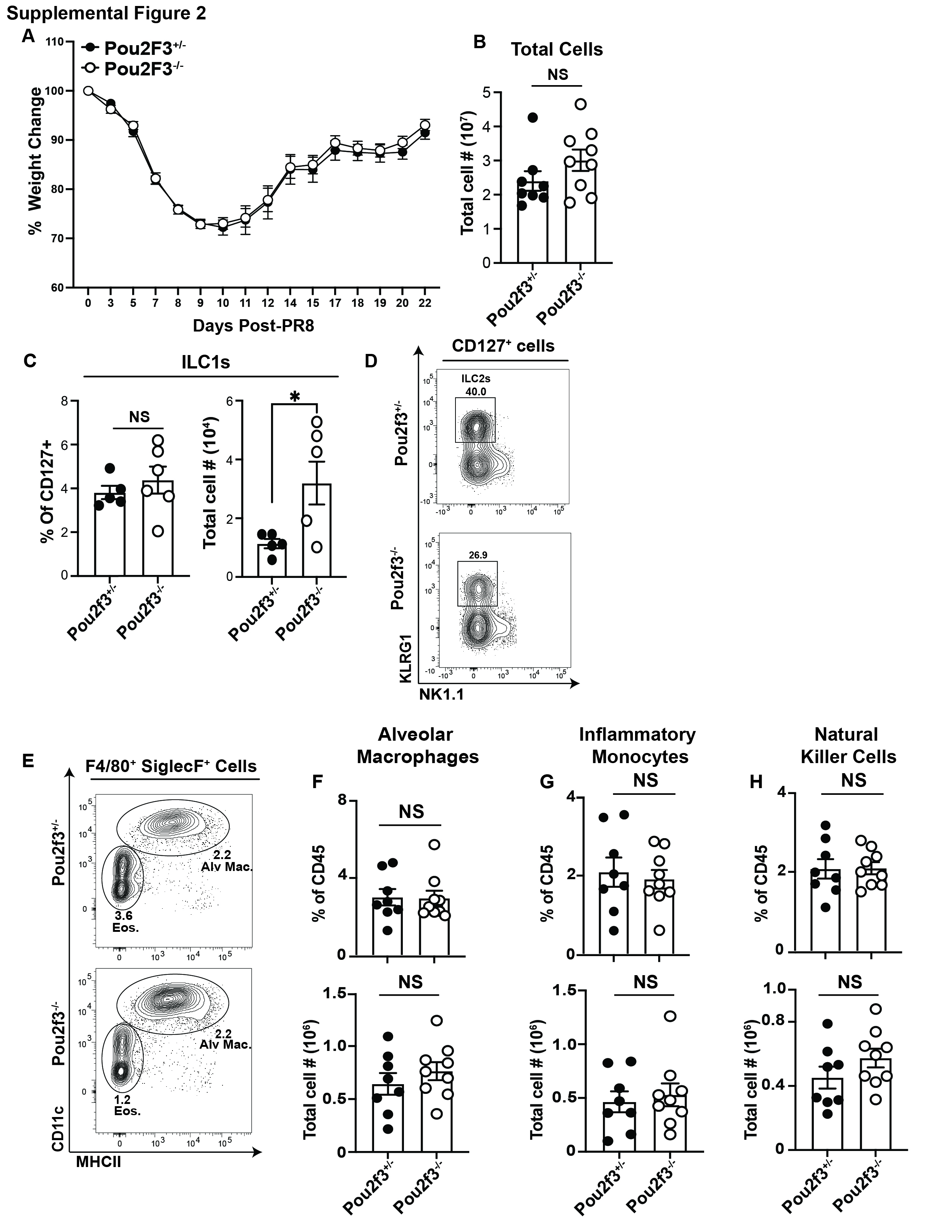

### Supplemental Figure 3

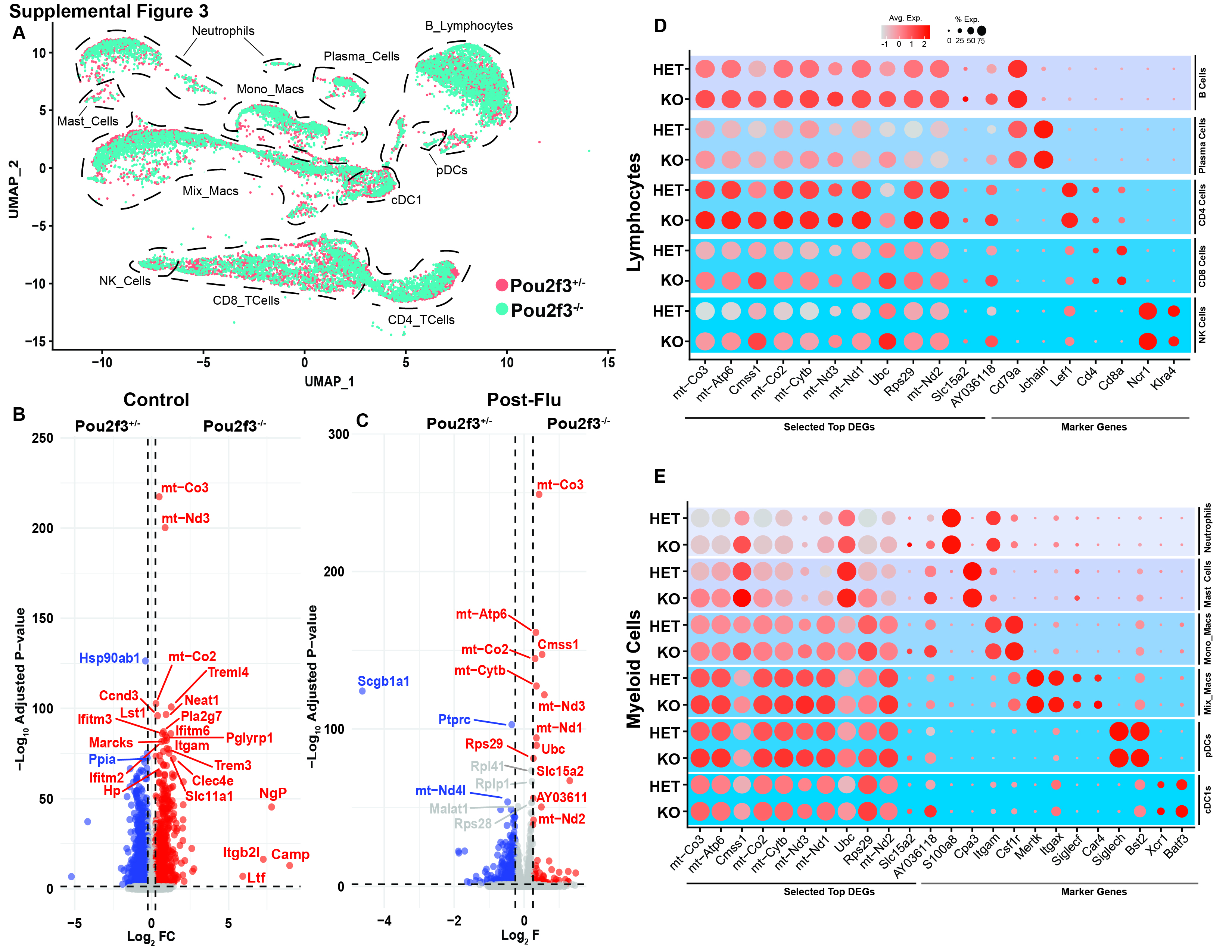

### Supplemental Figure 4

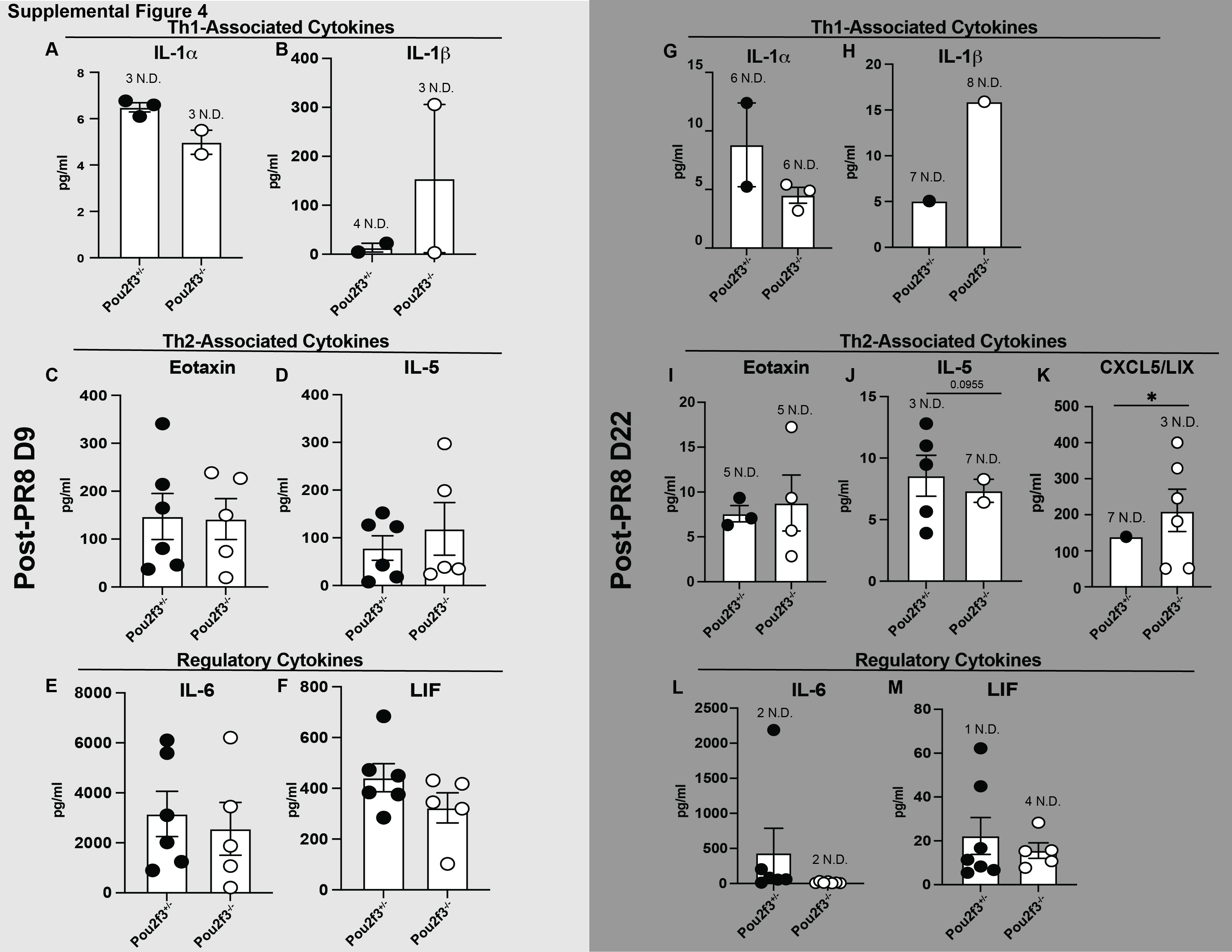

### Supplemental Figure 5

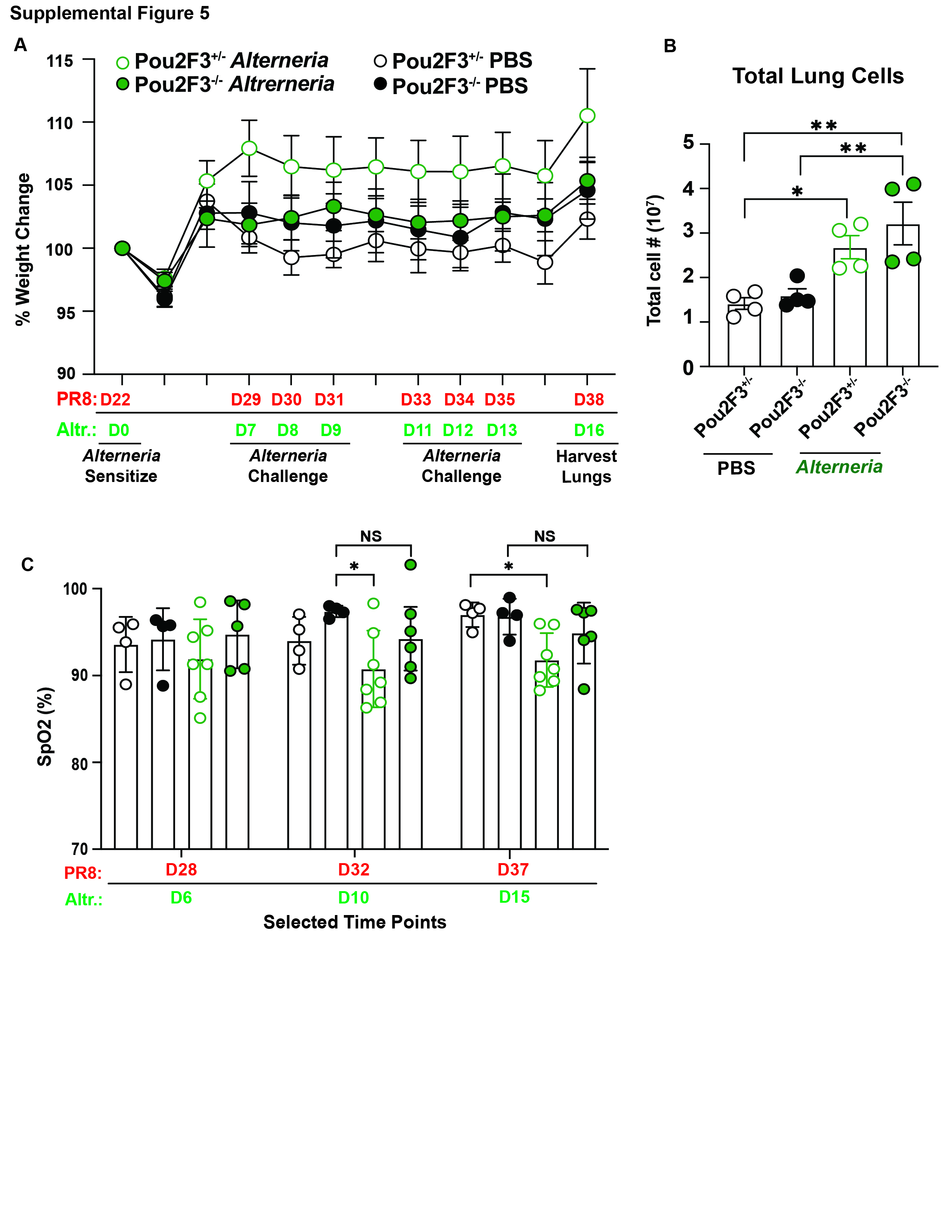

### Supplemental Figure 6

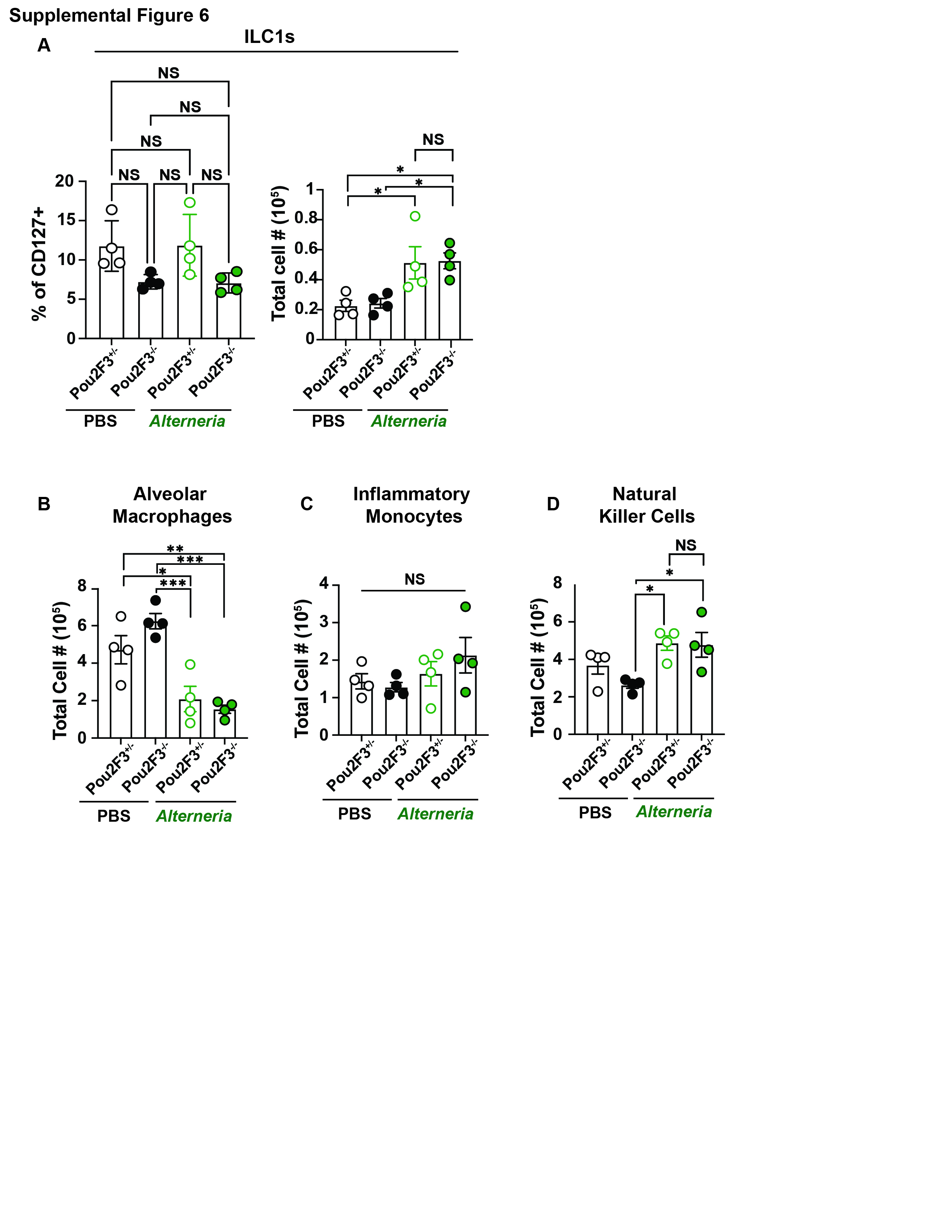
