## Supplemental Figure Legends for "Influenza-induced tuft cell expansion alters ILC-mediated inflammation"

**Supplemental Figure 1: Changes in body weight and oxyhemoglobin saturations following PR8 infection.**

Body weight and pulse oximetry were assessed prior to infection and at indicated time points following PR8 infection. **A-B**) WT (BL6) (n=2-5) and IFNγ^-/-^(n=3-8) mice, **C-D**) ILC1^+/-^ (n=3), and ILC1^-/-^ (n=3) mice and **E-F**) ILC2^+/-^ (n=3-5) and WT (n=3-5) mice. **A** and **E** combined two independent experiments. **B, D** and **F** each circle represents an individual mouse. **E)** Mice that lost <15% of starting body weight after influenza infection were excluded. Unpaired, two-tailed parametric t test was used at each time point to determine significance between knockouts and controls. Error bars = SEM

**Supplemental Figure 2: Lung innate immune cells at day 22 post PR8-infection.**

**A**) Changes in body weight throughout the course of infection in Pou2f3^+/-^ (n=5) and Pou2f3^-/-^ (n=6) mice. **B**) Total lung cell numbers of Pou2f3^+/-^ and Pou2f3^-/-^ mice at 22 days (D22) following PR8-infection (p.i.). **C**) Frequency of CD127^+^ and total cell numbers of ILC1s (Rorgt^-^Tbet^+^ of KLRG1^-^CD127^+^cells) at D22 p.i. in Pou2f3^+/-^ and Pou2f3^-/-^ mice. **D**) Representative contour plots of lung ILC2s (KLRG1^+^NK1.1^-^ of CD127^+^ Lin^-^ cells) at D22 p.i. in Pou2f3^+/-^ and Pou2f3^-/-^ mice. **E**) Representative contour plots of lung eosinophils (Eos: CD11c^low/int^ MHCII^-^ of SiglecF^+^F4/80^+^ cells) and alveolar macrophages (Alv. Mac.: CD11c^high^ MHCII^+^ of SiglecF^+^F4/80^+^ cells) in the lungs of Pou2f3^+/-^ and Pou2f3^-/-^ mice at D22 p.i. **F**) Frequency of CD45^+^ and total cell numbers of alveolar macrophages (CD11c^high^ MHCII^+^ of SiglecF^+^F4/80^+^ cells), **G**) inflammatory monocytes (Ly6c^high^CD11b^+^ of Ly6g^-^ cells) and **H**) natural killer cells (CD127^-^NK1.1^+^ of Lin^-^CD11b^low/int^ cells) in the lungs of Pou2f3^+/-^ and Pou2f3^-/-^ mice at D22 p.i. **B** and **F-H** represent combined two independent experiments. Each circle represents an individual mouse . P values were calculated using unpaired, two-tailed parametric t test at each time point between the genotypes (* = p < 0.05, NS = non-significant). Error bars = SEM. Complete gating strategy for the different immune cell populations found in material and methods.

**Supplemental Figure 3: Transcriptional profiling of immune cell populations in tuft cell-deficient mice post-influenza.**

**A**) Single-cell RNA-seq UMAP clustering of sorted immune (CD45^+^) cell types from mice lungs at D22 p.i. **B**) Volcano plot comparing gene expression in uninfected Pou2f3^+/-^ and Pou2f3^-/-^ mice **C**) Volcano plot comparing gene expression in infected Pou2f3^+/-^ and Pou2f3^-/-^ mice lungs at D22 p.i. Top 12 differentially expressed genes (DEGs) and selected marker genes in **D**) lymphocytes and **E**) myeloid cells in Pou2f3^+/−^ and Pou2f3^−/−^ mice lungs at D22 p.i.

**Supplemental Figure 4: Luminex® cytokine array of bronchoalveolar lavage fluid collected from Pou2f3^−/−^ mice and controls at days 9 and 22 post–PR8 infection.**

Protein concentration of cytokines collected from the bronchoalveolar lavage fluid (BALF) of Pou2f3^+/-^ and Pou2f3^-/-^ mice at D9 and D22 p.i. **A–F)** Protein concentrations of IL-1α, IL-1β, eotaxin, IL-5, IL-6, and LIF at D9 p.i. in the BALF of Pou2f3^+/−^ (n = 6) and Pou2f3^−/−^ (n = 5) mice. **G-M)** Protein concentrations of IL-1α, IL-1β, eotaxin, IL-5, CXCL5/LIX, IL-6, and LIF at D22 p.i in the BALF of Pou2f3^+/−^ (n = 8) and Pou2f3^−/−^ (n = 9) mice. **G-M** combined two independent experiments. Each circle represents an individual mouse. P values were calculated using unpaired, two-tailed parametric t test (* = p < 0.05, NS = non-significant). Statistical testing utilized a value of zero when protein concentration was below the limit of detection. Error bars = SEM.

**Supplemental Figure 5:** **Changes in body weight and oxyhemoglobin saturations following PR8 infection and subsequent allergen challenge.**

**A**) Body weight was measured at indicated time points following PR8 infection and *Alternaria alternata* challenge. Mice were initially infected with PR8 and allowed to recover, then sensitized on D22 with *A. alternata*  (40 µg) or vehicle control (PBS), followed by challenge on days 29, 30, 31, 33, 34, and 35 with *A. alternata* (20 µg) or PBS and harvested at D38 post-PR8. **B**) Total lung cell numbers in Pou2f3^+/−^ and Pou2f3^−/−^ mice at D38 p.i. following challenge with PBS or *A. alternata*; n= 4/group. **C**) Pulse oximetry were assessed at indicated time points following PR8 infection challenge with PBS or *A. alternata*. **A** and **C**) Pou2f3^+/−^ PBS (n = 4), Pou2f3^−/−^ PBS (n = 4), Pou2f3^+/−^ *A. alternata* (n = 7), and Pou2f3^−/−^ *A. alternata* (n = 6). P values were calculated using one-way ANOVA with Tukey’s post test for multiple comparisons for each time point collected (* p < 0.05. ** = p < 0.01. NS = non-significant). Error bars = SEM

**Supplemental Figure 6:** **Lung innate immune cells following PR8 infection and subsequent *Alternaria alternata* challenge.**

**A**) Frequency of CD127^+^ and total cell numbers of ILC1s (NK1.1^+^KLRG1^-^ of CD127^+^Lin^-^ cells) in Pou2f3^+/−^ and Pou2f3^−/−^ mice at D38 p.i. following challenge with PBS or *A. alternata*. **B**) Total numbers of alveolar macrophages (CD11c^+^MHCII^+^ of SiglecF^+^CD11B^int/high^ cells), **C**) inflammatory monocytes (Ly6c^high^CD11b^+^ of Ly6g^-^ cells) and **D**) natural killer cells (NK1.1^+^ CD127^-^ of Lin^-^ cells) in Pou2f3^+/−^ and Pou2f3^−/−^ mice at D38 p.i. following challenge with PBS or *A. alternata*. Each circle represents an individual mouse. P values were calculated using one-way ANOVA with Tukey’s post test for multiple comparisons. (* p < 0.05, ** p < 0.01, *** p < 0.001. NS = non-significant). Error bars = SEM. Complete gating strategy for the different immune cell populations found in material and methods.
